# Graphlet Laplacians: graphlet-based neighbourhoods highlight topology-function and topology-disease relationships

**DOI:** 10.1101/460964

**Authors:** Sam F. L. Windels, Noël Malod-Dognin, Nataša Pržulj

**Affiliations:** Department of Computer Science, University College London, London WC1E 6BT, UK

## Abstract

**Motivation:** Laplacian matrices capture the global structure of networks and are widely used to study biological networks. However, the local structure of the network around a node can also capture biological information. Local wiring patterns are typically quantified by counting how often a node touches different graphlets (small, connected, induced sub-graphs). Currently available graphlet-based methods do not consider whether nodes are in the same network neighbourhood.

**Contribution:** To combine graphlet-based topological information and membership of nodes to the same network neighbourhood, we generalize the Laplacian to the Graphlet Laplacian, by considering a pair of nodes to be ‘adjacent’ if they simultaneously touch a given graphlet.

**Results:** We utilize Graphlet Laplacians to generalize spectral embedding, spectral clustering and network diffusion. Applying our generalization of spectral clustering to model networks and biological networks shows that Graphlet Laplacians capture different local topology corresponding to the underlying graphlet. In biological networks, clusters obtained by using different Graphlet Laplacians capture complementary sets of biological functions. By diffusing pan-cancer gene mutation scores based on different Graphlet Laplacians, we find complementary sets of cancer driver genes. Hence, we demonstrate that Graphlet Laplacians capture topology-function and topology-disease relationships in biological networks

## 1 Introduction

Systems biology is flooded with large scale “omics” data. Genomic, proteomic, interactomic, metabolomic and other data, are typically modeled as networks (also called graphs). This abundance of networked data started the fields of network biology, allowing us to uncover molecular mechanisms of a broad range of diseases, such as rare Mendelian disorders (Smedley *et al.*, 2014), cancer (Leiserson *et al.*, 2015), and metabolic diseases (Chen *et al.*, 2008). In personalized medicine, network analysis is applied to the tasks of bio-marker discovery (Chen *et al.*, 2013), patient stratification (Hofree *et al.*, 2013) and drug repurposing (Yamanishi *et al.*, 2010).

Many network analysis methods use the Laplacian matrix, as it captures the global structure of a graph (see section 1.1). These methods include spectral clustering, spectral embedding and network diffusion. Each of these families of methods relies on the fact that the eigen-decomposition of the Laplacian matrix naturally uncovers network clusters (see section 1.2). Applications of spectral embedding include visualizing genetic ancestry (Lee *et al.*, 2010) and pseudo-temporal ordering of single-cell RNA-seq profiles (Campbell *et al.*, 2015). Applications of spectral clustering include detection of functional sub-network modules in single-cell genomic networks (Bartlett et al., 2017), construction of protein backbone fragments libraries (Cribben and Yu, 2017), and connectivity analyses and computational modeling of human brain function from fMRI data (Craddock *et al.*, 2012). Network diffusion methods are widely used for protein function prediction (e.g. (Cao *et al.*, 2013)) and discovery of disease genes and disease modules (e.g. (Leiserson *et al.*, 2015)), see (Cowen *et al.*, 2017) for a full review.

### 1.1 Laplacian matrix definition

The *Laplacian matrix* captures the global structure of a network: for each node it captures the adjacency relationship with other nodes and its degree centrality. In a network, *G*(*V, E*), two nodes, *u* and *v*, are *adjacent* if there exists an edge (*u, v*) ∈ *E* connecting them. The adjacency of all nodes in graph *G* is represented in an *n* × *n* symmetric *adjacency matrix A*:

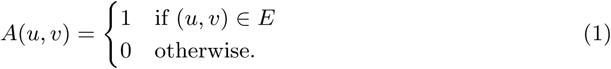

The *neighbourhood* of a node is defined as the set of nodes adjacent to it. A node’s *degree* is the size of its neighbourhood, or equivalently, the number of nodes that are adjacent to it. The *degree matrix* of *G* is defined as the diagonal matrix, *D*, where *D*(*u, u*) is equal to the degree of node *u*:

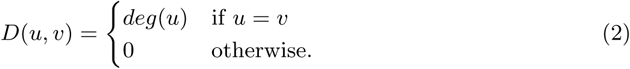

The Laplacian, *ℒ*, is defined as *ℒ* = *D* − *A*. The symmetrically normalized Laplacian, *L^sym^*, is defined as *L^sym^* = *D*^−1/2^*ℒD*^−1/2^.

### 1.2 Laplacian matrix eigendecomposition

Spectral clustering, spectral embedding and laplacian based network diffusion, analyze networks based on the eigenvectors of the Laplacian matrix, which naturally uncover clusters present in the network as they are the solution to (an approximation of) the ratio-cut problem. The *eigenvectors* of *ℒ* are the set of vectors such that their linear transformation by *ℒ* corresponds to the same vector rescaled by a scalar called the *eigenvalue*. They are found as the result of the *eigendecomposition* of *ℒ*, which solves *ℒU* = *U*Λ, where the *i*-th column of the *n × n* matrix *U* is the *i*-th eigenvector, *u_i_*, of *ℒ*, and Λ is the diagonal matrix whose diagonal elements are the corresponding eigenvalues. The *ratio-cut problem* aims to cut a graph into *d* similar-sized graph partitions whilst minimizing the number of edges being cut. An approximation of this problem is formulated as follows:

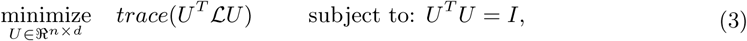

where each column of *U* is a normalized indicator vector assigning each node to one of the *d* graph partitions. This problem is solved by *d* normalized eigenvectors of *ℒ* associated with the *d* smallest eigenvalues, illustrating how they capture clusters present in the network.

### 1.3 Matrix alternatives to the Laplacian

Laplacian matrices only capture direct interactions between nodes. To capture the influence of long-range interactions between nodes, Estrada (2012) proposed the *k-path Laplacian* by generalizing the concepts of adjacency and degree. The *k*-path Laplacian defines a pair of nodes *u* and *v* to be *k-adjacent* if the shortest path distance between them is equal to *k*. Analogously, *k-path degree*, deg_*k*_(*u*), generalizes the concept of the degree to the number of length *k* shortest paths that have node *u* as an endpoint. The *k*-path laplacian, 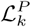, is defined as:

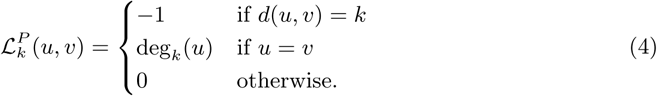

Because of biotechnological capturing limitations and human biases, biological networks are globally noisy and locally less noisy, so methods that are designed to capture the local structure of the data have been shown to outperform global methods (Roweis and Saul, 2000; Wu and Scholkopf, 2006).*Vicus* is an alternative to the Laplacian that captures the intricacies of a network’s local structure (Wang *et al.*, 2017) based on the semi-supervised network label propagation algorithm of Zhou *et al.* (2004). Label propagation is defined as *P* = *BQ*, where the *n* × *d* matrix Q assigns the *n* nodes of network *G* to one of *d* possible labels (if the node is labeled), *B* is an *n* × *n* diffusion matrix (see further for details), and the reconstructed matrix *P* is an *n* × *d* matrix used for predicting labels for unlabeled nodes. By assuming that *P* ≈ *Q* and setting Vicus, *ℒ^V^*, to be equal to (*I* − *B^T^*)(*I* − *B*), it was shown that *Q* can be learned as the eigenvectors of *ℒ^V^*. Vicus captures the label diffusion, *Q* captures the local connectivity between nodes that is implied given the diffusion, leading to the interpretation of *ℒ^V^* as a Laplacian matrix.

The label diffusion matrix, *B*, is defined as follows. The authors of Vicus modified the diffusion process slightly from the original, in the sense that diffusion is limited to the direct neighborhood of each node to give Vicus its ‘local’ characteristics (instead of the diffusion process being applied over the entire graph at once). Formally, they define for each node *u* a local *K* × *K* adjacency matrix *W_u_*, which is the sub-matrix of the adjacency matrix *A* limited to *u* and its *K* − 1 neighbours, *N*(*u*). Then, *S_u_* is defined as the row-normalized transition matrix of *W_u_*:

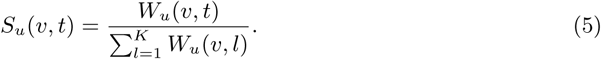

Matrix *β_u_* encodes the label diffusion for *u* and its direct neighbours *β_u_* = (1 − α)(I − *αS_u_*)^−1^, where α controls the level of label diffusion. Label diffusion matrix, *B*, of *G* is defined as:

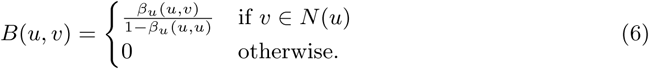

Vicus has been applied to protein module discovery and ranking of genes for cancer subtyping (Wang *et al.*, 2017).

### 1.4 Problem

The *k*-path Laplacian captures the influence of long-range interactions between nodes, but ignores short-range interactions. Although Vicus is focused on capturing local topology, it lacks interpretability from a structural perspective. Graphlet based methods have shown that in protein-protein interaction (PPI) networks the local wiring patterns contain biological information. Graphlets are small, connected sub-graphs (see section 2.1 for a formal definition) used to capture the local topology around a node in a network. They have been applied to predict protein function (Milenković and Pržulj, 2008; Davis *et al.*, 2015) and to identify new cancer genes (Milenković *et al.*, 2010) directly from the similarities in terms of their interaction patterns in PPI networks. However, as current graphlet-based methods summarize local topology by a vector of counts, they lose the information of whether the nodes being compared are in the same neighbourhood.

### 1.5 Contribution

We introduce the Graphlet Laplacian, allowing us to analyze nodes based on their network neighbourhoods, whilst restricting the pattern of their interactions to that of a prespecified graphlet. Hence, each graphlet (Figure 1.A) has its own corresponding Graphlet Laplacian. We generalize spectral embedding, spectral clustering and network diffusion to utilize Graphlet Laplacians. Through graphlet-generalized spectral clustering of model networks and biological networks, we show that different Graphlet Laplacians capture different local topology. By applying graphlet-generalized spectral embedding, we visually demonstrate that Graphlet Laplacians capture biological functions as well. We quantify this through graphlet-generalized spectral clustering analysis. We show that Graphlet Laplacians are not only as biologically relevant as alternative Laplacian matrices, but also capture complementary biological functions. Finally, by graphlet-generalized diffusing of pan-cancer gene mutation scores on the human PPI network, we show that Graphlet Laplacians capture complementary disease modules.

**Figure 1:**
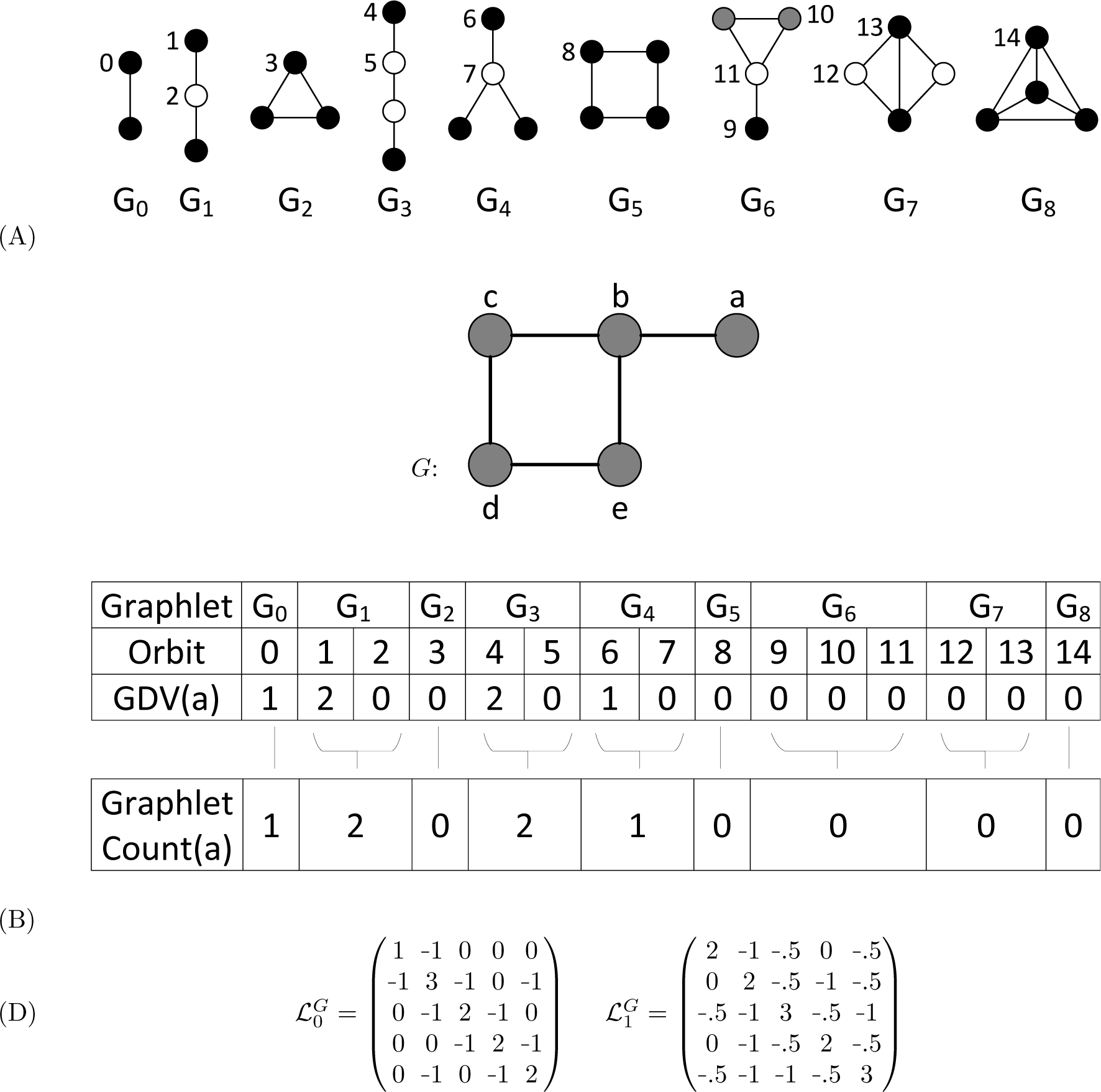
Illustration Graphlets and Graphlet Laplacians. **A:** All graphlets with up to 4 nodes, labeled *G*_0_ to *G*_8_. The 15 automorphism orbits are colored and labeled within each graphlet. **B:** The graphlet degree vector of node ‘a’ in the example network, *G*, and its relationship to graphlet counts. Node ‘a’ touches graphlet *G*_0_ once on orbit 0, via edge a-b. Node ‘a’ touches graphlet *G*_1_ twice, each time at orbit 1, via paths a-b-c and a-b-e. **C:** The Graphlet Laplacians for graphlets *G*_0_ and *G*_1_, applied on the network, *G*, shown in panel B. The diagonal elements correspond to the graphlet counts of each node; e.g. 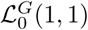 is equal to 1, the number of times node ‘a’ touches graphlet *G*_0_, 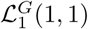 is equal to 2, the number of times node ‘a’ touches graphlet *G*_1_. The off-diagonal elements correspond to the number of times two nodes touch a given graphlet together, scaled by *size* (*G_k_*) − 1. 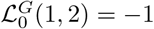, as ‘a’ and ‘b’ form *G*_0_ once and *size*(*G*_0_) − 1 = 1. 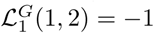, as ‘a’ and ‘b’ form *G*_1_ twice and *size* (*G*_1_) − 1 = 2.

## 2 Materials and methods

### 2.1 Graphlet Laplacian definition

We generalize the concept of the Laplacian to that of a Graphlet Laplacian by generalizing the definitions of the adjacency and degree to the ones based on graphlets.*Graphlets* are small, connected, non-isomorphic, induced sub-graphs of a large network (Pržulj *et al.*, 2004) and are used to capture the local topology around a node in a network. Within each graphlet, automorphism orbits are groups of nodes that can be swapped (Pržulj, 2007). The *graphlet degree vector* (GDV) of a node contains the counts of how many times a node touches each graphlet at a particular automorphism orbit (Milenković and Pržulj, 2008). It is used to quantify the local topology around a node. Figure 1.B. illustrates the GDV of node *a* in the example network and its relationship in graphlet counts. That is for a given node, the graphlet count is equal to the sum of the orbit counts corresponding to that graphlet. We define two nodes *u* and *v* of *G* to be *graphlet-adjacent* with respect to a given graphlet, *G_k_*, if they simultaneously touch *G_k_*. Note that for *k* ∈ {1, 3, 4, 5, 6, 7}, *u* and *v* can be graphlet-adjacent without there being an edge (*u, v*) between them. For example, in Figure 1.B, nodes *b* and *d* are graphlet-adjacent with respect to graphlet *G*_5_, despite (*b, d*) not being an edge. By this definition of adjacency, it is possible for two nodes to be adjacent multiple times, as there can exist different subsets of nodes, *S*, by which the two nodes touch graphlet *G_k_*. Therefore, we can measure the strength of the relationship between two nodes with respect to graphlet *G_k_*, by counting how often they simultaneously touch *G_k_*:

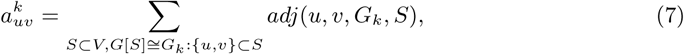

where *adj*(*u, v, G_k_, S*) is equal to 1 if *u* and *v* simultaneously touch graphlet *G_k_* that is induced by set *S* of nodes of *G*, and 0 otherwise. Given this extended definition of adjacency, we define the graphlet based adjacency matrix as:

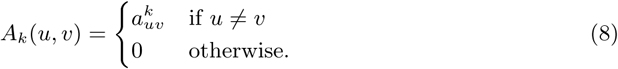

Analogously, the *graphlet degree* generalizes the node degree as the number of times node *u* touches graphlet *G_k_*:

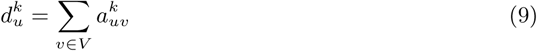

Similarly, we extend the degree matrix to the *Graphlet Degree matrix* for graphlet *G_k_*, *D_k_*:

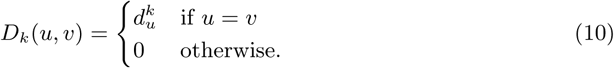

For an underlying graphlet *G_k_*, we define the *Graphlet Laplacian*, 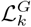, as:

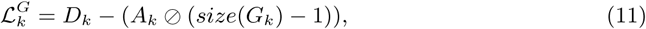

where ⌀ represents the element-wise division operator and *size*(*G_k_*) is the number of nodes of *G_k_*. 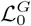 and 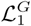 are illustrated in Figure 1.C.

### 2.2 Graphlet Laplacian properties

To allow for an easy interpretation of the Graphlet Laplacian for each graphlet, *G_k_*, we introduce the two-step transformation function, *T*, which maps graph *G* to its Graphlet Laplacian representation: 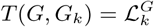. First, *T* converts *G* = {*V, E*} to a weighted network *G*′ = {*V, E*′}, where the weight of each edge (*u, v*) in *G*′ corresponds to 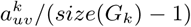 measured in *G*. Next, *T* converts *G*′ to its standard Laplacian representation. This shows that the Graphlet Laplacian can be interpreted as the Laplacian of an undirected weighted network. Therefore, our Graphlet Laplacian retains the following key properties of the Laplacian (Mohar *et al.*, 1991; Merris, 1994):

1. The Graphlet Laplacian, 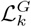, is symmetric and positive semi-definite.
2. The smallest eigenvalue is 0 and the corresponding eigenvector is the constant vector **1**.
3. The Graphlet Laplacian has *n* non-negative, real-valued eigenvalues: 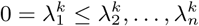.
4. The multiplicity of the eigenvalue 0 equals the number of connected components in *G*′, which we refer to as *graphlet based components*.

Finally, note that the Graphlet Laplacian for graphlet *G*_0_, 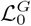, is equivalent to the standard Laplacian, *ℒ*.

### 2.3 Spectral embedding

Spectral Embedding embeds a network in a lower dimensional space, placing nodes close in space if they share many neighbours. Here, we generalize the Laplacian Eigenmap embedding algorithm (Belkin and Niyogi, 2003) so that two nodes are embedded close in space if they frequently simultaneously touch a given graphlet. Given an unweighted network *G* with *n* nodes, we find a low dimensional embedding, *Y* = [**y**_1_,…,**y**_*n*_] ∈ ℝ^*d×n*^ such that if nodes *u* and *v* are frequently graphlet-adjacent with respect to graphlet *G_k_*, then **y**(*u*) and **y**(*v*) are close in the *d*-dimensional space by solving:

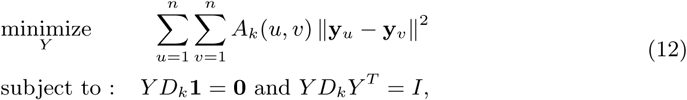

where *A_k_* is the graphlet-based adjacency matrix of *G* for graphlet *G_k_*, *D_k_* is the graphlet-based degree matrix of *G* for graphlet *G_k_*. The columns of *Y* are found as the generalized eigenvectors associated with the second to (d+1) smallest generalized eigenvalues solving 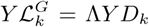, where Λ is the diagonal matrix with the generalized eigenvalues along its diagonal.

### 2.4 Spectral clustering

Spectral clustering groups nodes in a network that form densely connected network clusters. By generalizing spectral clustering to Graphlet Laplacian based spectral clustering, we are able to identify network components that are similarly wired with respect to a given graphlet. Many different variations of spectral clustering exist (Von Luxburg, 2007). Aiming for a balanced clustering, we generalize normalized spectral clustering as defined by Ng *et al.* (2002) to use different Laplacians including Graphlet Laplacians, all denoted by a generic *ℒ* in algorithm 1. We skip the normalization step (i.e. step 1) for Vicus, as Vicus is already normalized.

#### Algorithm 1 Normalized spectral clustering

**Input** Number of clusters, *d*, and Laplacian matrix, *ℒ*, of an unweighted network, *G*, with *n* nodes.

**Output** *d* clusters of the *n* nodes of *G*.

1: Compute the normalized laplacian as:*ℒ^sym^* = *D*^−1/2^*ℒD*^−1/2^.

2: Compute the *d* eigenvectors of *ℒ^sym^*. associated with its *d* smallest eigenvalues: *Y* = [**y**_1_,…,**y**_*d*_] ∈ ℝ^*n×d*^.

3: Next, we normalize *Y* so that each row has unit norm.

4: Cluster the *n* points 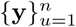 into *d* groups using k-means.

For each network, we determine the numbers of clusters, *d*, by using the rule of thumb: 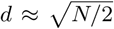 (Kodinariya and Makwana, 2013). In the Supplement Section 1, we present the justification for this approach, based on inspection of the spectra of different Laplacian matrices of each network. Because of the heuristic nature of spectral clustering, we perform 20 runs for each clustering and consolidate them into a single clustering applying ensemble clustering (Ghosh and Strehl, 2002).

### 2.5 2.5 Network diffusion

Network diffusion refers to a family of related techniques, which propagate information on nodes through the network. Here, we will focus on generalizing the diffusion kernel to *graphlet based diffusion kernel*. The diffusion kernel is often called the ‘heat kernel’, as it can be viewed as describing the flow of heat originating from the nodes across the edges of a graph with time. In network biology, nodes typically represent genes and ‘heat’ on a node represents experimental measurements. For a set of *n* nodes, these measurements are encoded in vector *P*_0_ ∈ ℝ^*n*^. Information is diffused as follows:*P* = *H P*_0_, where *H* is a diffusion kernel. For a given graphlet *G_k_*, we define the graphlet based diffusion kernel, 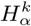, as:

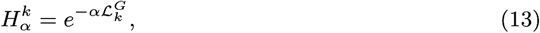

where the parameter α ∈ ℝ controls the level of diffusion. This way, diffusion of information on nodes propagates between nodes restrained by how often they form a given graphlet *G_k_* together.

### 2.6 Topological dissimilarity of networks

The Graphlet Correlation Distance (GCD-11) is the current state of the art heuristic for measuring the topological distance between networks (Yaveroğlu *et al.*, 2014). First, the global wiring pattern of a network is captured in its Graphlet Correlation Matrix (GCM), an 11 × 11 symmetric matrix comprising the pairwise Spearman’s correlations between the 11 non-redundant orbit counts over all nodes in the network. The Graphlet Correlation Distance between two networks is computed as the Euclidean distance of the upper triangle values of their GCMs.

### 2.7 Cluster enrichment analysis

To assess if a cluster of genes is biologically relevant, we measure if it is statistically significantly enriched in a specific biological annotation term by applying the hyper-geometric test. That is, we consider each cluster as a’sampling without replacement’, in which each time we find a given annotation, we count that as a’success’. The probability of observing the same or higher enrichment (i.e. successes) of the given annotation purely by chance is equal to:

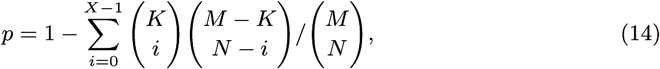

where *N* is the number of annotated genes in the cluster, X is the number of genes annotated with the given annotation in the cluster, *M* is the number of annotated genes in the network, and *K* is the number of genes annotated with the given annotation in the network. An annotation is considered to be statistically significantly enriched if its enrichment p-value, after correction for multiple hypothesis testing (Benjamini and Hochberg, 1995), is lower than or equal to 5%.

### 2.8 Data

#### 2.8.1 Real biological network data collection

We create three types of molecular interaction networks for human and baker’s yeast (*S. cerevisiae*) by collecting the following data: experimentally validated protein-protein interactions (PPIs) from IID version 2018-05 (Kotlyar *et al.*, 2016) and BioGRID version 3.4.161 (Stark *et al.*, 2006), genetic interactions from the same version of BioGRID, and gene co-expressions from COXPRESdb version 6.0 (Okamura *et al.*, 2015).

#### 2.8.2 Random model network generation

We generate ten networks containing 2,000 nodes at edge density of 1.5% (to mimic our biological networks), for each of the following widely used seven random network models: Erdös-Rènyi random graphs (ER) (Erdös Paul and Rényi Alfréd, 1959), generalized random graphs with the degree distribution matching to the input graph (ER-DD) (Newman, 2010), Barabási-Albert scale-free networks (SF-BA) (Barabási and Albert, 1999), scale-free networks that model gene duplication and mutation events (SF-GD) (Vazquez *et al.*, 2001), geometric random graphs (GEO) (Penrose, 2003), geometric graphs that model gene duplications and mutations (GEOGD) (Pržulj *et al.*, 2010), and stickiness-index based networks (Sticky) (Pržulj and Higham, 2006). As our real biological networks have power-law degree distributions, our set of model networks contains four types of networks with power-law degree distribution: ER-DD, SFBA, SF-GD and Sticky. The GEO and GEO-GD random network models are generated using 3-dimensional space.

#### 2.8.3 Biological annotations

For each gene in our biological networks, we collect the most specific experimentally validated biological process annotations (BP), cellular component annotations (CC) and molecular function annotations (MF) present in the Gene Ontology (GO) (Ashburner *et al.*, 2000; Gene Ontology Consortium, 2017).

#### 2.8.4 Cancer gene annotations

We collect the pan-cancer gene mutation frequency scores computed by Leiserson *et al.* (2015) for the purpose of detecting of pan-cancer disease modules. Leiserson *et al.* (2015) collected raw pan-cancer mutation data, such as SNV’s, indels and CNA’s, from the TCGA database (Kandoth *et al.*, 2013). These data were filtered to exclude statistical outliers and include only the samples (corresponding to a patient) for which SNV and CNA data were available. The resulting data set contains mutations on 11,565 genes across 3,110 patients in cancers across 20 different tissues. Additionally, we collect the sets of known cancer driver genes in all available tissues from IntOGen (Gonzalez-Perez *et al.*, 2013) and Cosmic (Futreal *et al.*, 2004).

## 3 Results and discussion

### 3.1 Graphlet Laplacians capture different local topology

We apply Graphlet Laplacian based spectral clustering for different underlying graphlets to partition a network into different sets of sub-networks. To decide if two Graphlet Laplacians capture different topology, we create a distribution of distances (measured by GCD-11, see section 2.6) between the sub-networks captured by each of them. To test the significance of this distance distribution, we compare it against a baseline, which is the distribution of distances between sub-networks of one of the two partitionings to itself. Statistical significance of this comparison is determined using the Mann-Whitney U test. Figure 2 illustrates this approach for the comparison of the topology captured by 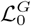 and 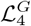 in the human PPI network. We observe that the distribution of distances between the sub-networks captured by 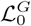 and 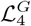 is statistically significant, with a p-value of 0.0. In general, we find that clusters obtained from different Graphlet Laplacians are typically statistically significantly topologically different at the 5% significance level. This is true across all of our biological networks and most of our model networks (with the exception of GEO), as shown in the Supplement, Section 2. This demonstrates that different Graphlet Laplacians capture different local topology in the same network.

**Figure 2:**
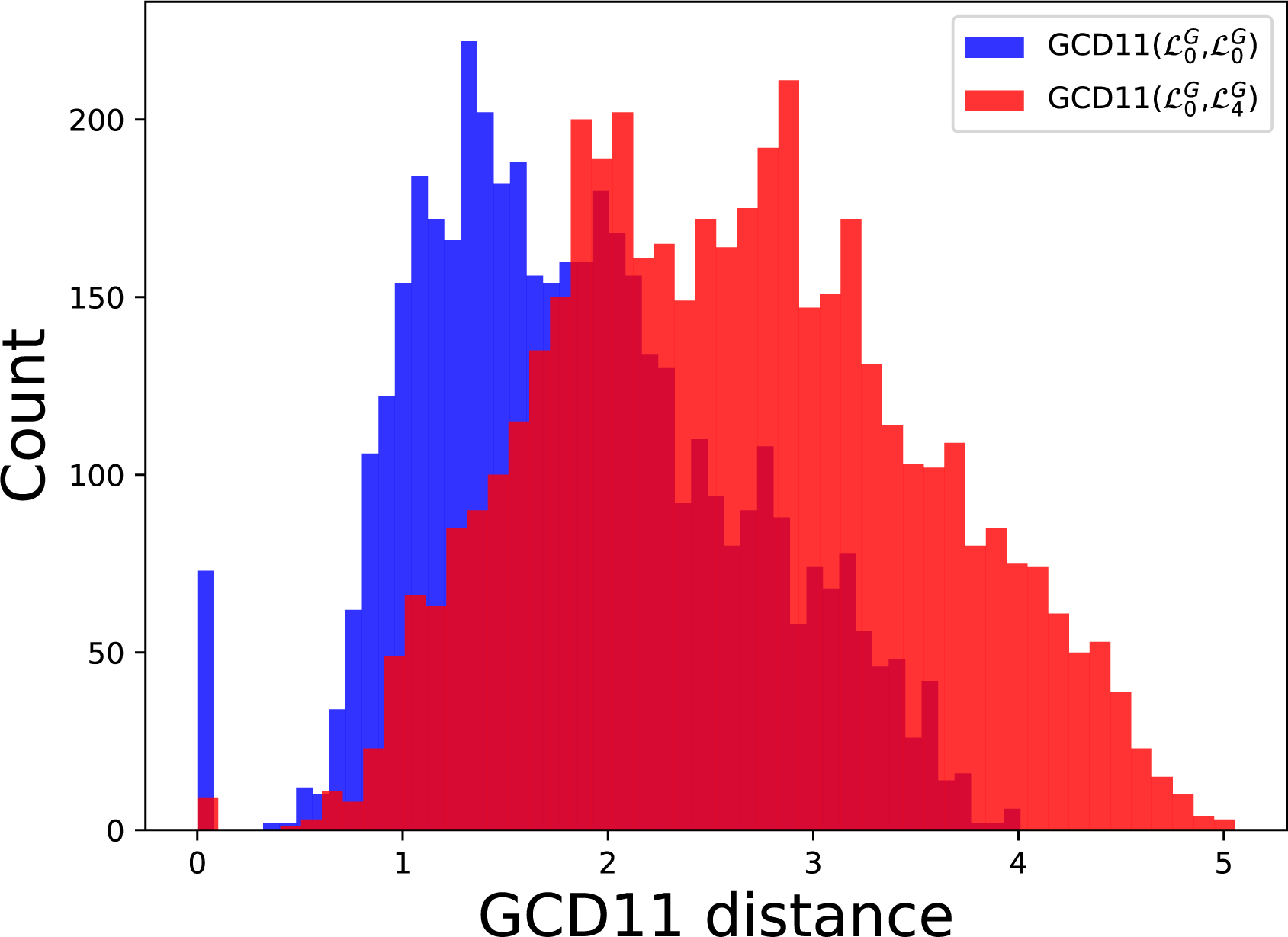
Comparison of topological distance distributions between sub-networks captured by two different Graphlet Laplacians in the human PPI network. We show the distribution of GCD-11 distance of the sub-networks captured by 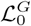 in the human PPI network to itself (blue), and the distribution of distances to those captured by 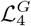 (red).

### 3.2 Different Graphlet Laplacians capture different biological function

In addition to showing that our Graphlet Laplacians capture different local topology, we assess their capacity to capture biological functions. To informally visualize this, we perform spectral embedding. We focus on the embedding of the yeast GI network, for which we use 14 core biological process annotations defined by Costanzo *et al.* (2016). We illustrate the spectral embedding of the symmetrically normalized 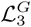 Graphlet Laplacian in Figure 3. The embeddings of the other Laplacian matrices of the yeast GI network can be found in the Supplement, Section 3.

**Figure 3:**
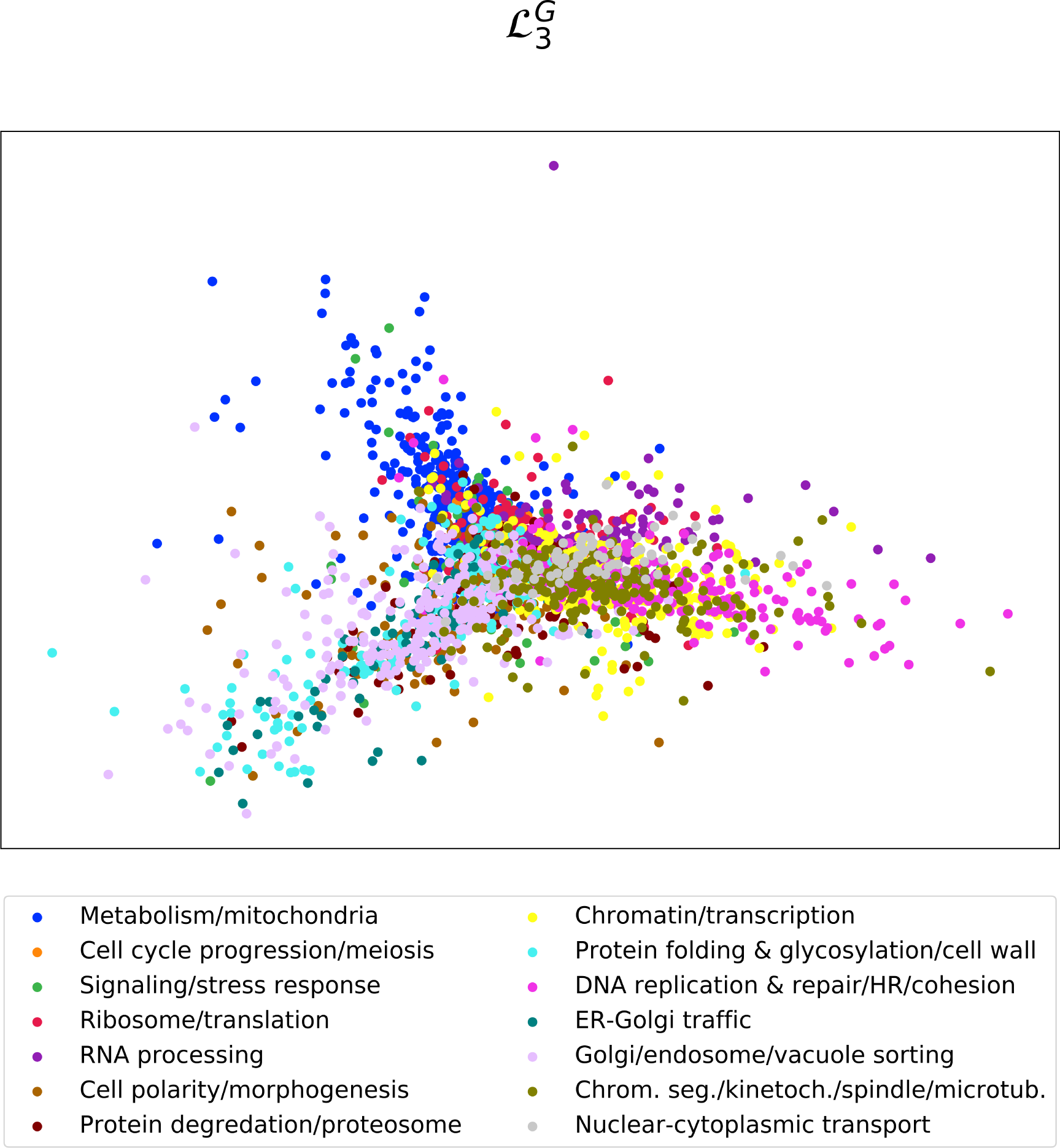
Capturing biological functions with Graphlet Laplacian 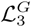. 2D spectral embedding of the yeast GI network using the Graphlet Laplacian for *G*_3_. Points represent genes and are color-coded with 14 core biological process annotations defined by Costanzo *et al.* (2016).

As seen in Figure 3, the spectral embedding of 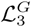 correctly groups and separates the biological processes of ‘nuclear cytoplasmic transport’, ‘metabolism / mitochondria’, ‘Golgi / endosome / vacuole sorting’ and ‘Chrom. seg. / kinetoch. / spindle / micro tub.’. In the supplement, we illustrate that Vicus and the Laplacian fail to find any grouping at all, placing all of the nodes in the same dense cluster. Embeddings based on 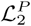 and 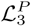 succeed in separating different genes into different clusters, but without grouping them in a biologically meaningful way.

Next, we quantify the functional separation captured by Graphlet Laplacians. We apply Graphlet Laplacian based spectral clustering for each graphlet on our set of human molecular networks and assess the functional enrichments in terms of the percentage of clusters enriched and the total number of annotations enriched (Figure 4). First, we observe that clusterings based on all Graphlet Laplacians but 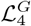 tend to be of similar quality as those based on the standard Laplacian or Vicus, both in terms of percentage of clusters enriched as well as total number of annotations enriched. 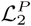 and 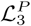 capture almost no function in PPI networks, as indicated by the low percentages of clusters enriched and relatively few enriched GO-BP annotations found. We find similar results in yeast, see Supplement, Section 5. In the Supplement, we additionally observe that for each network and annotation type, there is always at least one Graphlet Laplacian that shows a larger number of the total number of enriched annotations than Vicus. We conclude that Graphlet Laplacians are at least as biologically as relevant as the standard Laplacian, *k*-path Laplacian and Vicus.

**Figure 4:**
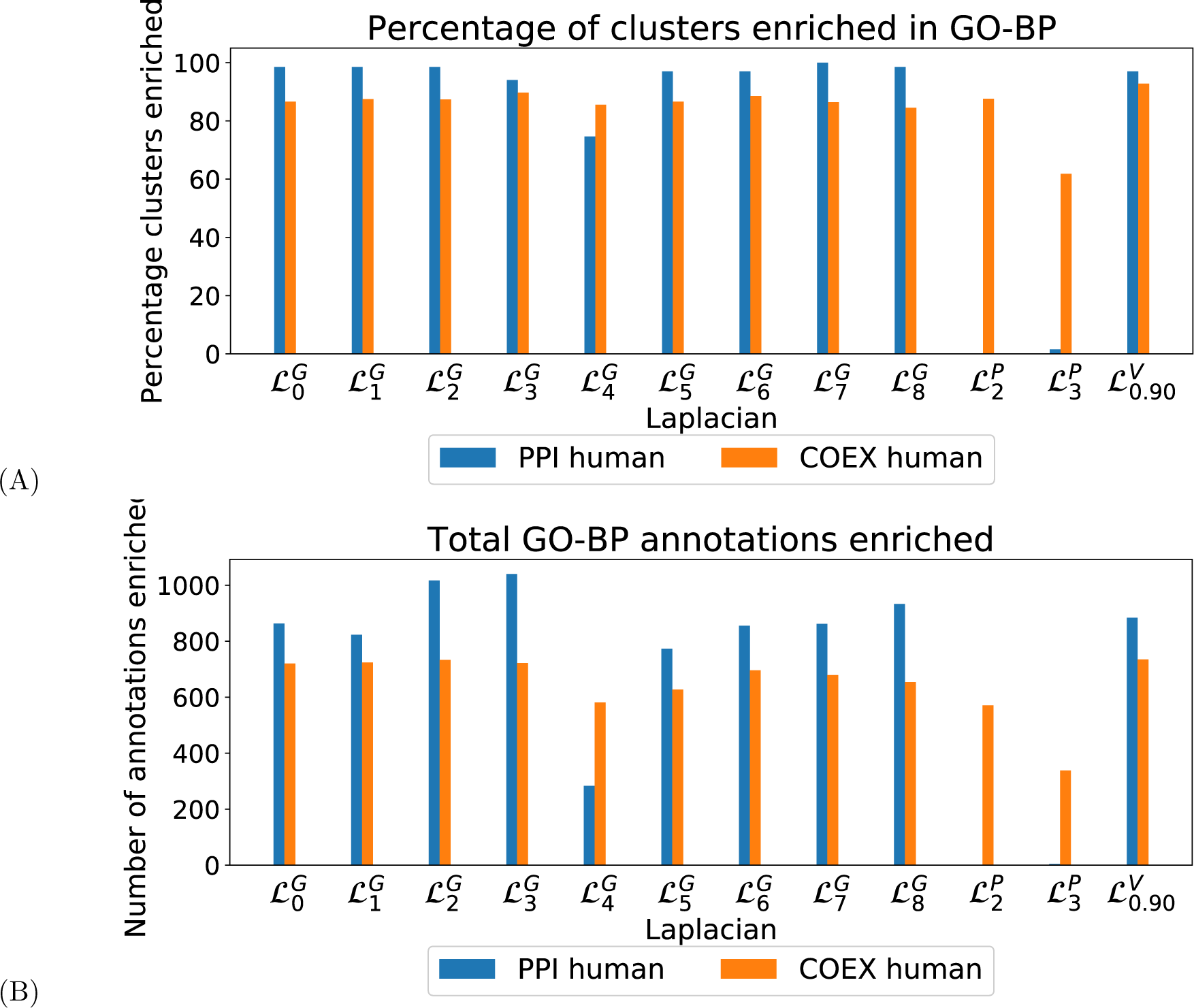
Cluster Quality. **A:** For our set of human molecular networks (color-coded), the percentage of clusters enriched in BP annotations, with clusters obtained based on spectral clustering using different Laplacian matrices (x-axis). **B:** For our set of human molecular networks, the total number of enriched GO-BP annotations in clusters obtained based on spectral clustering using different Laplacian matrices (x-axis).

Having established that Graphlet Laplacian based clusters capture biological functions, we quantify the overlap in their enriched functions. In the Supplement, Section 6, we calculate the Jacard Index between the sets of enriched functions corresponding to each Graphlet Laplacian. For GO-BP enrichments in clusterings on the human PPI and COEX networks, the average Jacard Index is 0.5 and 0.55 respectively, meaning that different Graphlet Laplacians capture different functions. To further demonstrate this point, we present the number of GO-BP functions that are enriched only in the clustering obtained by a particular Graphlet Laplacian (Figure 5). We observe that each type of Laplacian matrix shows a tendency to capture some distinct biological functions, indicating the link between the biological function and the topology of these diverse molecular networks. We The same is observed for GO-MF and GO-CC annotations, both in yeast and human networks (see the Supplement, Section 7). Combining this observation with our previous results, we can conclude that Graphlet Laplacian based spectral clustering allows for distinguishing different sets of similarly wired network components that are not only biologically relevant, but may also capture complementary biological functions.

**Figure 5:**
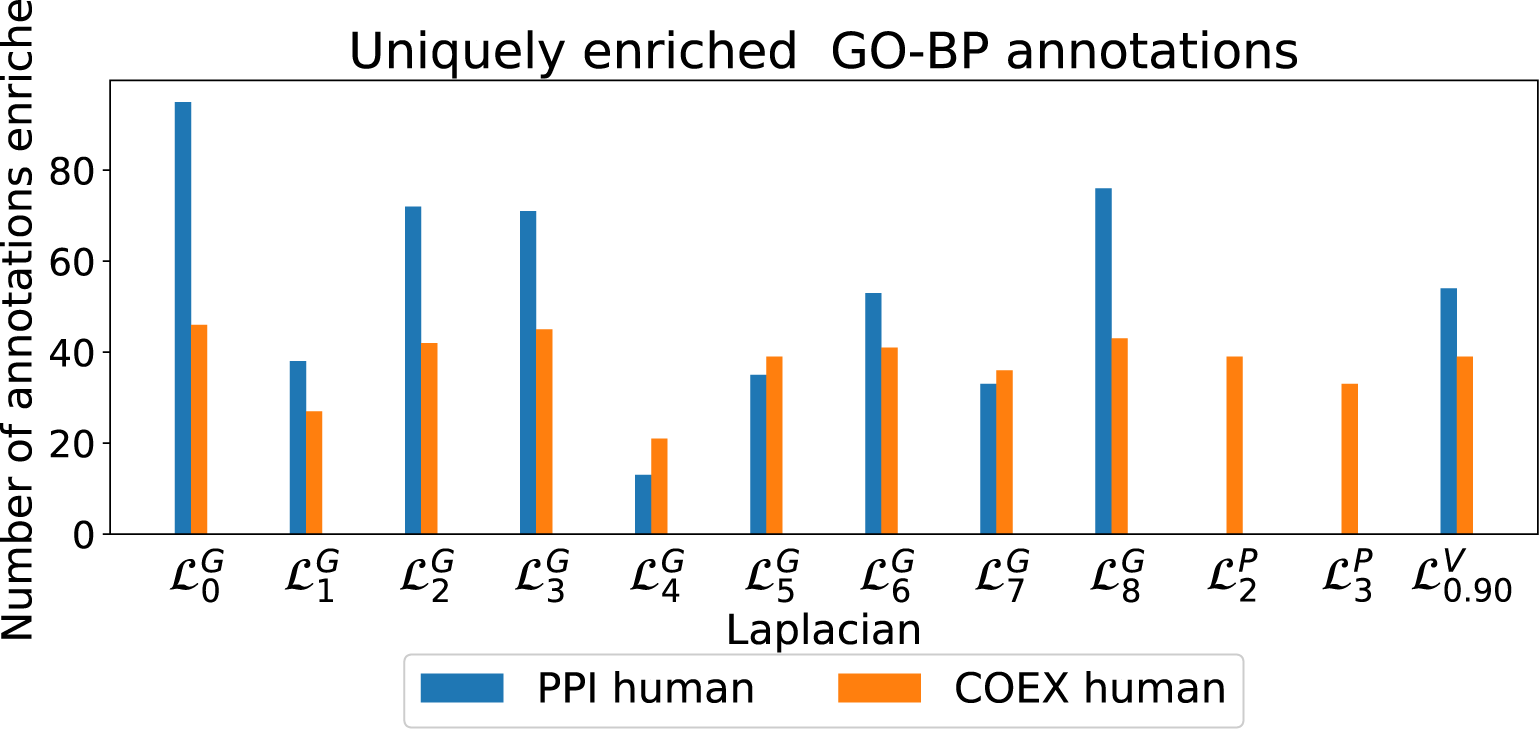
GO-BP uniquely enriched. The number of annotations that are uniquely enriched in clusterings based on the indicated Laplacian matrix for each biological network (color coded).

### 3.3 Different Graphlet Laplacians predict complementary sets of pan-cancer driver genes

Finally, we show that Graphlet Laplacians can be used to predict complementary sets of cancer driver genes. We do this by diffusing (see section 2.5) the gene mutation frequency scores (see section 2.8.4) on the human PPI network based on different Graphlet Laplacian matrices. For each graphlet laplacian, we rank the diffused scores and use them as cancer driver gene predictions. We assume a gene is correctly predicted as a cancer driver gene if it is known to be a cancer driver gene in at least one type of cancer (see section 2.8.4). We measure the accuracy of our predictions using the area under the Precision-Recall (PR) curve and the area under the Receiver Operator Characteristic (ROC) curve. In Supplement Section 7, we show the prediction accuracy for the diffusion of the gene mutation frequency scores using different Graphlet Laplacians. We observe that prediction accuracy is independent of the Graphlet Laplacian used and on par with the standard Laplacian, with an average area under the PR and ROC curve of 0.21 and 0.78 respectively. In terms of accuracy Graphlet Laplacian based predictions consistently outperform predictions based on *k*-path or Vicus, which achieve an average area under the PR and ROC curve of 0.17 and 0.74 and 0.14 and 0.73 respectively. In figure 6 we evaluate the overlap between the top hundred cancer driver gene predicted per Laplacian, measured using the Jacard Index. We observe five distinct clusters of different Laplacian matrices predicting different sets of cancer driver genes. Importantly, diffusion based on three sets of Graphlet Laplacians (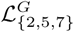, 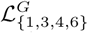 and 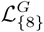) provide predictions dissimilar to those achieved using the standard Laplacian (the average Jacard Index of each cluster with the standard Laplacian based predictions being 0.75, 0.80, 0.65 respectively). Conversely, the highest scoring genes based on 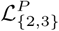 prove to overlap greatly with those based on the standard laplacian (the average Jacard Index being 0.95). Vicus based diffusion provides cancer driver gene scores dissimilar from all other laplacian matrices, be it at lower accuracy, as shown above. Similar results are obtained applying graphlet generalized diffusion on the human COEX network, as shown in Supplement, Sections 8 and 9. We conclude that Graphlet Laplacian based diffusion can be used to find complementary sets of cancer driver genes.

**Figure 6:**
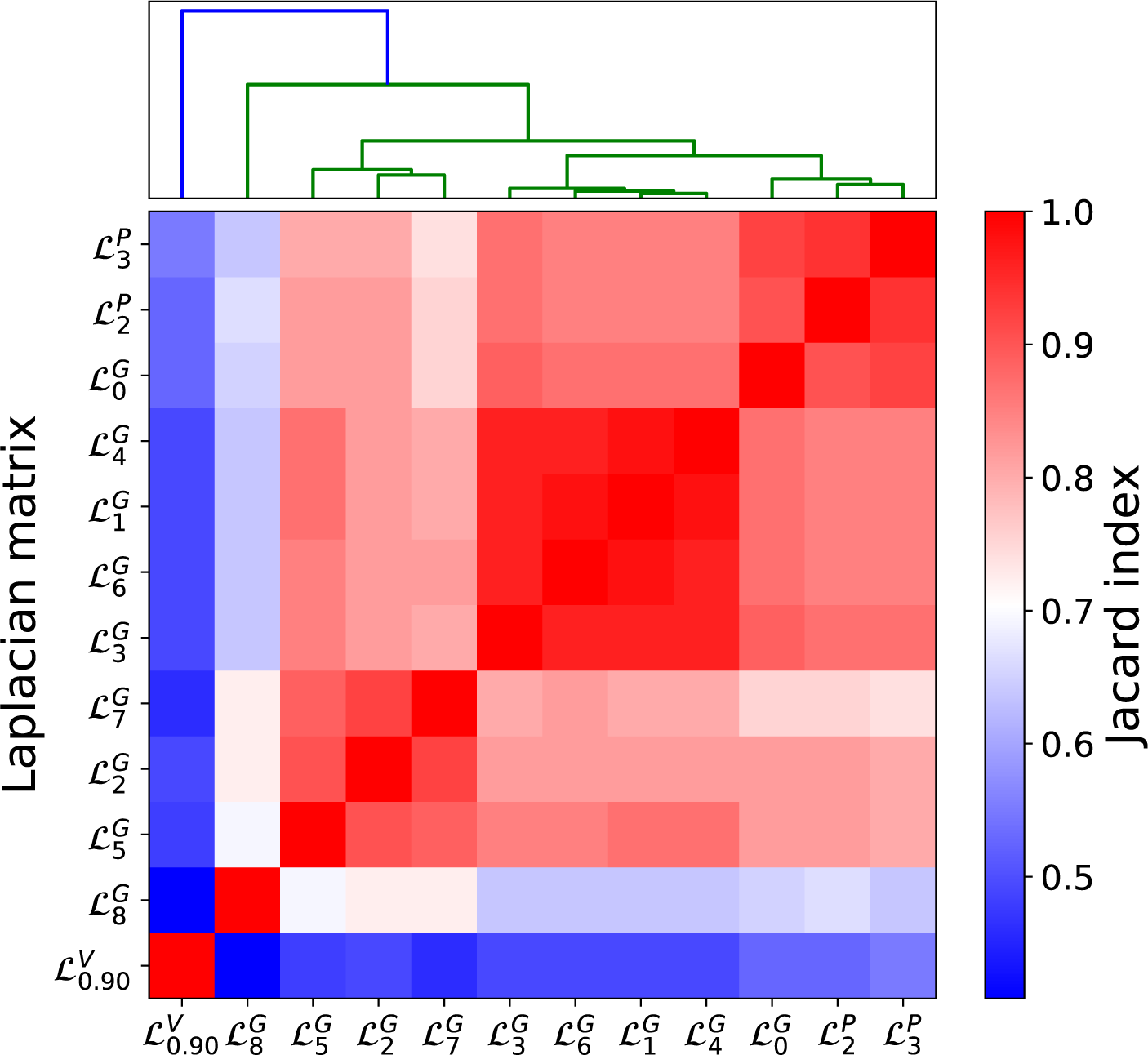
Overlap of highest scoring cancer driver genes. Evaluating the overlap in the sets of the top 100 predicted genes based on different Laplacians measured by the Jacard Index, with predictions computed by performing network diffusion of mutation frequency scores on the human PPI network.

## 4 Conclusion

To combine graphlet-based topological information and membership of nodes in the same network neighborhood, we generalize the Laplacian to the Graphlet Laplacian, by considering two nodes to be graphlet-adjacent if they simultaneously touch a given graphlet. Through our generalized spectral clustering of model networks and biological networks, we showed that different Graphlet Laplacians capture sub-networks of distinct local topology. Furthermore, the sub-networks captured by Graphlet Laplacians in molecular networks are enriched in different and complementary sets of biological annotations. Although Vicus also achieves enrichments different from the standard Laplacian at a similar rate, results are not as readily interpretable from a topological perspective as they are with Graphlet Laplacians and the *k*-path Laplacian. With respect to the *k*-path Laplacian, Graphlet Laplacians cover a larger set of different local wiring patterns that could capture biological meaning. In PPI networks for example, the *k*-path Laplacian is not capable to capture biological signal in the sense of enriched GO-BP annotations in the resulting. Finally, we showed that our generalized network diffusion of pan-cancer gene mutation scores resulted in complementary sets of cancer driver genes dependent on the underlying graphlet. The accuracy of Graphlet Laplacian based predictions slightly outperform Vicus.*k*-path Laplacian based diffusion on the human PPI network added no additional driver gene information compared to the standard Laplacian.

In this paper, we introduced Graphlet Laplacians and demonstrated that they can straightforwardly be plugged into current Laplacian based network analysis methods widely used in systems biology, using spectral clustering, spectral embedding and network diffusion as example applications. However, Graphlet Laplacians could be used to leverage a whole range of the existing applications in Systems Biology and Network Medicine whose methods include the Laplacian. For example, Graphlet Laplacians could be used for the inclusion of prior knowledge in data integration methods for patient stratification (Hofree *et al.*, 2013; Gligorijević *et al.*, 2016), drug-target interaction prediction (Gu *et al.*, 2016), drug-drug interaction prediction (Zhang *et al.*, 2017). They could also be used for functional connectivity analysis in the brain (Abdelnour *et al.*, 2014) and spectral based network alignment (Stuart *et al.*, 2003).

